## Supplementary Material for "Context-Dependent Interaction Between Goal-Directed and Habitual Control Under Time Pressure"

September 28, 2024

### I. Methods

#### i. The Action Sequence Task

##### i.1 Criterion Test and Test Trials

After the instructions and before the main experiment, participants were told that their understanding of the task will be tested by a short test. In the test, participants were presented 13 successive dual-target trials. Participants were explicitly told that there is no time-limit during these test trials, and that they should always choose the option with the higher reward probability when possible. Participants were further told the stimulus positions with the higher reward probabilities. If a participant chose a low-reward-probability option in more than one of the 13 trials, they were again instructed about the test and their task. If failed, participants could repeat the test a maximum of two times, resulting in a maximum of three possible iterations. If a participant failed all three iterations, they were excluded from further participation. After successful completion of the criterion test, participants performed 20 test trials to get used to the pace of the experiment.

##### i.2 Counterbalancing

To account for hemispheric effects as well as for learning effects, participants were randomly assigned to one of four different counterbalancing groups, as shown in table S1.
**Mirroring.** Mirrored counterbalancing groups were introduced to account for hemispheric effects, like for instance a preference for the right hand (which is the dominant hand for the majority of the population). In reward contingency 1, the stimulus positions at the top left and bottom right had reward probabilities of 80%, while bottom left and top right had 20%. These were reversed in reward contingency 2, essentially mirroring the rewards along the vertical. Sequence 2 was a mirrored version of sequence 1, mirrored along the vertical, so if a stimulus appeared at the top left in sequence 1, it appeared at the top right in sequence 2. The stimuli in dual-target trials were mirrored in the same way for counterbalancing. Mirroring both stimuli and rewards ensured that congruent (incongruent) trials remained congruent (incongruent) even after mirroring.
**Reversed block orders.** Reversal of block orders was done to account for learning effects (Table

S1). Half of participants started with the random condition on day one and the repeating condition on day two, while the other half started with the repeating condition on day one and the random condition on day two. If all participants started for instance with the random condition on day one, reduced reaction times and error rates in the repeating condition (see main manuscript) could not be distinguished from possible learning effects at the beginning of the experiment.

#### i.3 Difference to previously published Task Version

The task paradigm used for this study was a revised and refined version of the previously published AST (Frölich et al., 2023). In the previously published version of the AST, some DTT trials featured two stimuli with the same reward probability (*Neutral DTTs*, with both stimuli either 20% or both 80% reward probability). For the present paradigm, we limited DTT types to random, congruent, and incongruent, to have one goal-directed response option in every DTT. Furthermore, in the previously published task version (Frölich et al., 2023), feedback consisted of a euro coin in the case of a point reward, and a white dot in the case of no point reward. To make feedback for both outcomes more similar, in the present version, feedback consists of a green smiley (point reward) and a red frowney (no point reward). The proportion of DTTs varied around 15% in the previous AST. In the present version, each block of 480 trials contains exactly 72 (15%) DTTs per block, with exactly 36 incongruent and 36 congruent DTTs in the repeating sequence condition. Lastly, in the previous version of the AST, stimulus sequences for the random condition were created pseudo-randomly subject to some constraints, and continuously for a whole block of 480 trials. In the present version, stimulus sequences for the random condition consisted of concatenated sequences of 12 elements, where each such sequence was created subject to the same constraints as the sequence for the Rep condition. Concatenation was performed such that no stimulus is repeated twice in a row.

#### i.4 Participants

Participants were randomly assigned to one of the 4 counterbalancing groups. Of 175 participants who initiated the experiment, 25 (14.3%) did not pass the Criterion Test. 3 participants timed out, 3 were excluded for initiating the study twice, 1 could not start part 2 of the experiment due to a user error, and 1 participant had technical problems during execution of the experiment. 10 participants could not finish the study due to a user error on the researchers' side, and 10 because of technical problems with the recruitment platform. 41 participants did not finish the experiment due to unknown reasons. 81 participants completed the experiment on both days. Of those 81 participants, 13 were excluded because they did not finish both parts of the experiment at approximately the same time of day or on two consecutive days, and 3 were excluded because of large error rates (error rates of $> 15\%$ in either single-target trials or dual-target trials or both). This resulted in 65 participants eligible for data analysis, (16 in counterbalancing group 1, 15 in group 2, 18 in group 3, 16 in group 4). In order for each counterbalancing group to contain the same number of participants for data analysis, we randomly excluded 1 participant from group 1, 3 from group 3, and 1 from group 4. This resulted in 15 participants in each of the four groups.

**Table S1:** The four different counterbalancing groups.

| | **Sequence 1 with Reward Contingency 1** | **Sequence 2 with Reward Contingency 2** |
|---|---|---|
| **Block Order 1** | Group 1 | Group 3 |
| **Block Order 2** | Group 2 | Group 4 |

**Table S2:** The two different block orders.

| | Block order 1 | Block order 2 |
|---|---|---|
| Day 1 | Rep | Rand |
| | Rand | Rep |
| | Rep | Rand |
| | Rand | Rep |
| | Rep | Rand |
| | Rand | Rep |
| Day 2 | Rand | Rep |
| | Rep | Rand |
| | Rand | Rep |
| | Rep | Rand |
| | Rand | Rep |
| | Rep | Rand |
| | Rand | Rep |
| | Rep | Rand |

#### i.5 Sequence Generation

Stimulus sequences were generated as sequences of 12 elements. Such 12-element sequences was generated in such a way that the same stimulus position did not appear twice in a row, and such that a certain stimulus position followed another stimulus position only once in the sequence (so for instance 1-2-1-2-... was inadmissible). Each 12-element sequence contained each of the four stimulus positions exactly three times. For each block (480 trials) of the Random-Sequence condition, 40 different 12-element sequences were randomly created and concatenated such that no repeating stimulus positions occurred. 72 Dual-target trials were inserted pseudo-randomly into each block of 480 trials, such that DTTs are separated by two STTs at least and nine STTs at most, and in a way that the distribution of joker types and separations between jokers were roughly uniformly distributed.

### ii. Generative Modeling

#### ii.1 Drift-Diffusion Modeling

**Model fitting.** For the drift-diffusion model, fitting was performed using the HSSM toolbox Fengler et al. (prep). Model estimation was done in Python (Version 3.11.9). For group-level means we used uniform priors defined over numerically plausible parameter ranges (see code and data availability section for details). Model-fitting was done using MCMC sampling. Chain convergence was assessed via the Gelman-Rubinstein convergence diagnostic $\hat{R}$ and sampling was continued until $1 \leq \hat{R} \leq 1.05$ for all group-level and individual-subject parameters. DDM was fitted with collapsing bounds to simulate time pressure, and lapse probabilities, that is, the probability of a random choice.

### II. Results

#### i. Positive Measures of Habit

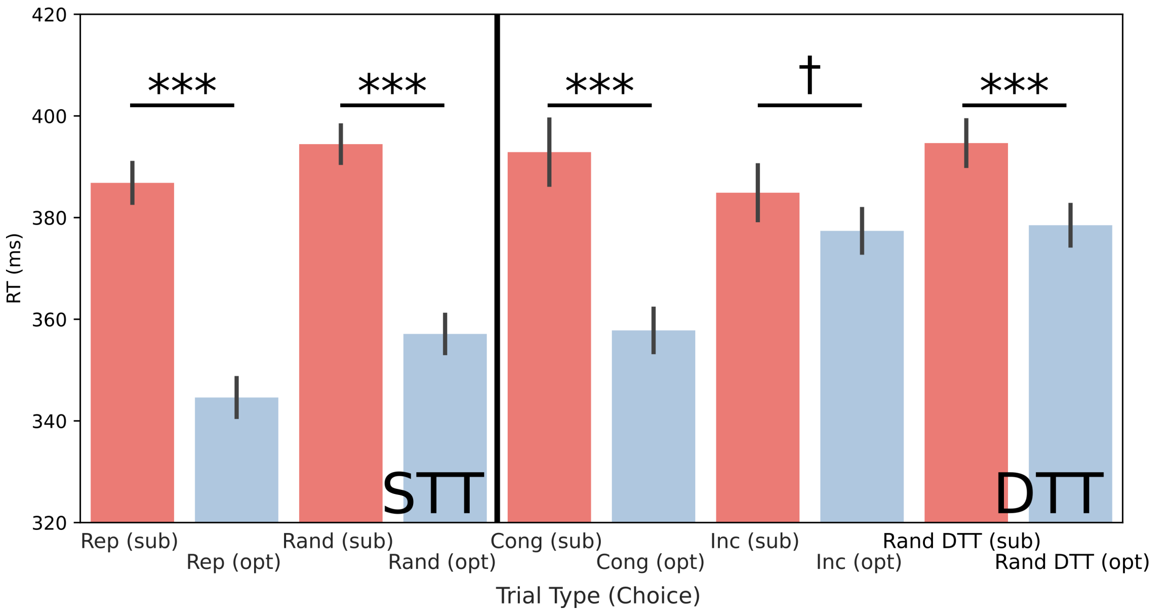


**Figure S1: Top**: Differences in optimal choices for participants with $\Delta$RT>0 (left) and $\Delta$RT 0 (right). **Bottom**: Differences in optimal choices for participants with $\Delta$ER>0 (left) and $\Delta$ER 0 (right).

#### ii. Generative Modeling and Model Comparison

Fig. S2 shows the posterior parameter distributions for the winning model (model 3). Fig. S3 shows the changes in posterior means of parameters from day 1 to day 2. Fig. S4 shows the differences of WAIC between models of the winning model family (Model Family 1) for each participant.

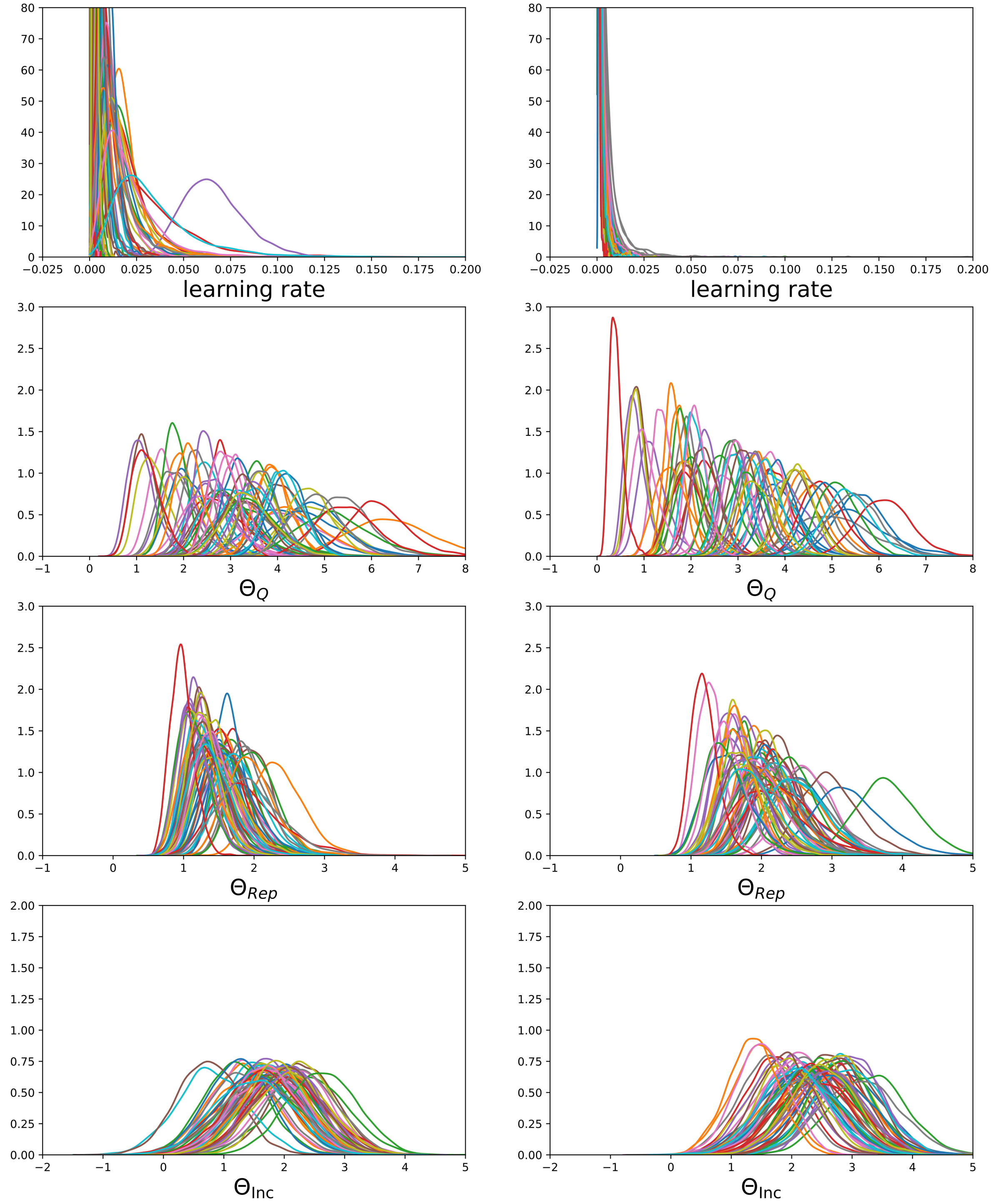


**Figure S2:** Posterior Distributions of inferred parameters of the best-performing model M 3.

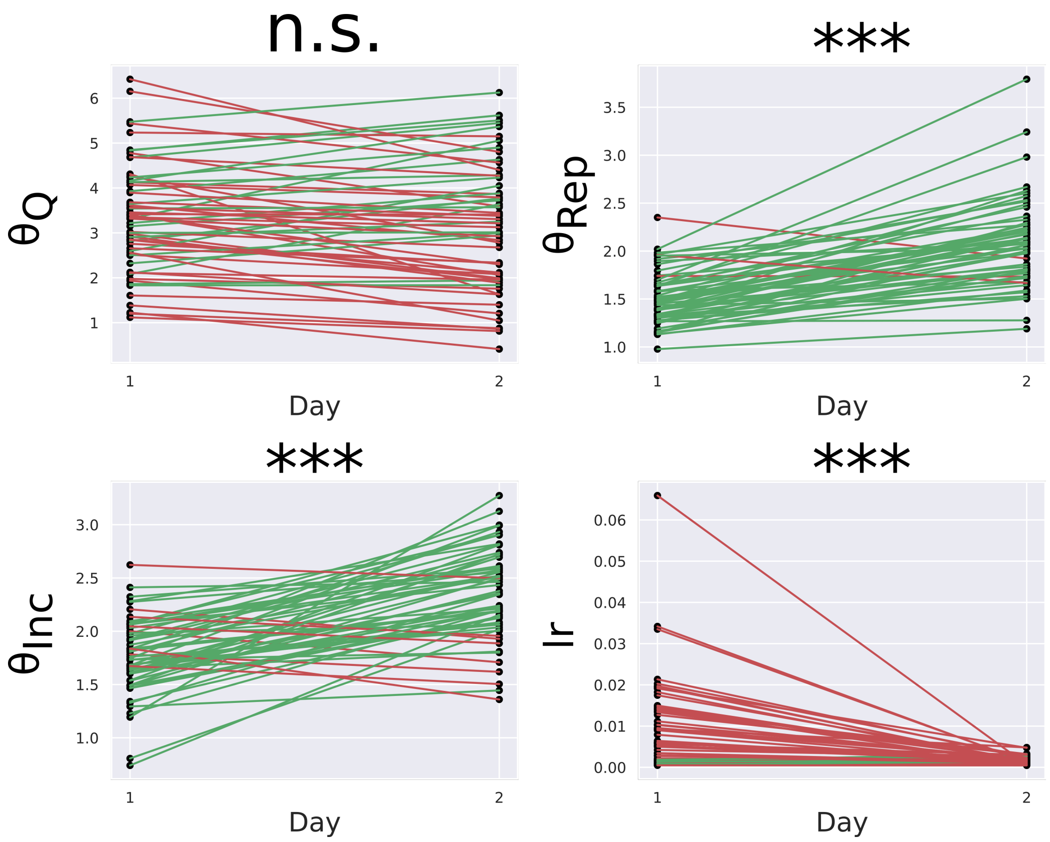


**Figure S3:** Changes of posterior means from day 1 to day 2. Learning rate, $\theta_{Rep}$, and $\theta_{Switch}$ increases significantly from day 1 to day 2 (paired two-sample t-tests). $*** : p < 0.001, ** : p < 0.01, * : p < 0.05$.

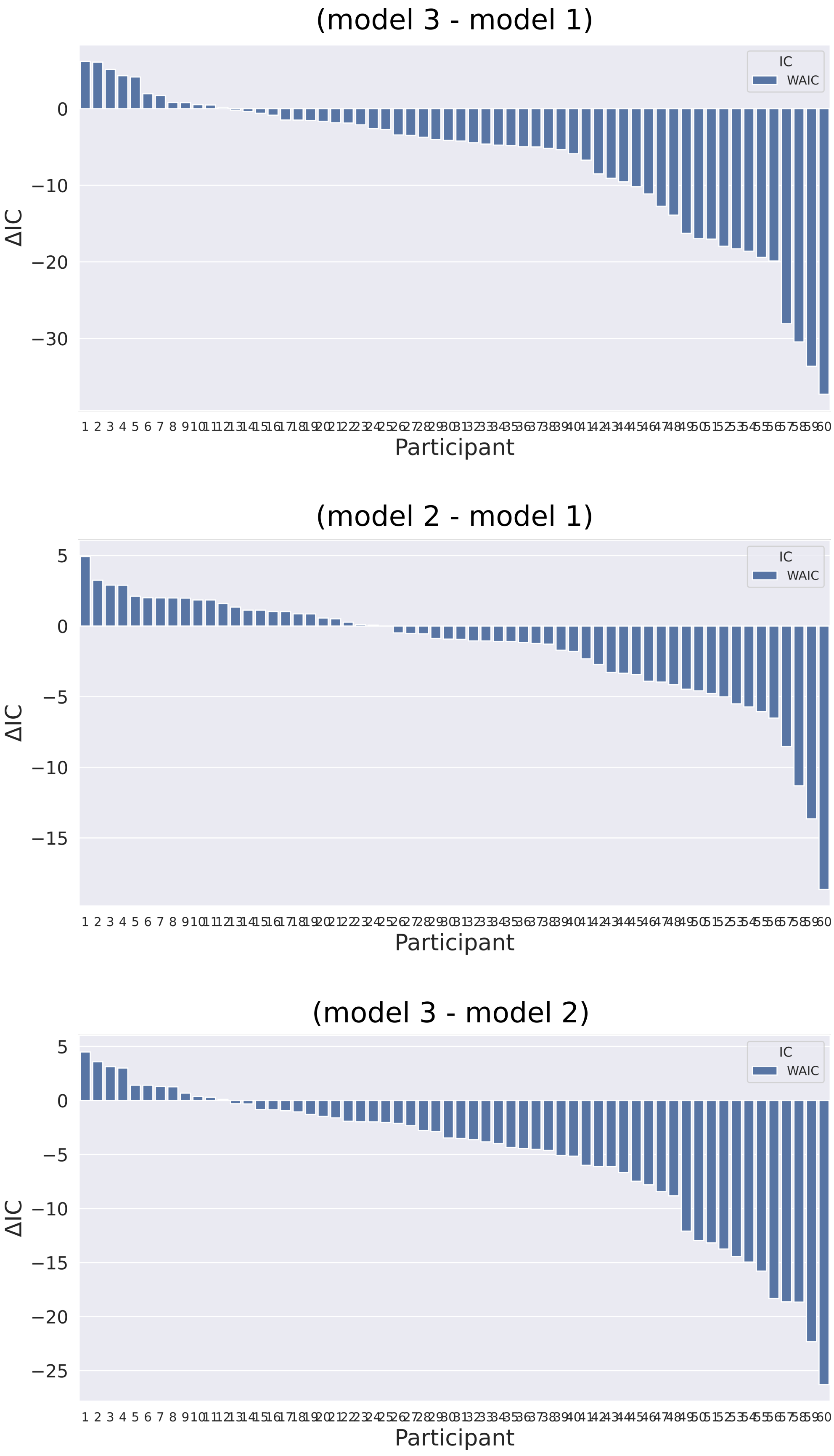




**Figure S4:** Differences of WAIC and DIC between model 1 and model 3 for each participant. Positive values indicate a preference for model 3.

### iii. Drift-Diffusion Modeling

Fig. S5 shows the distributions of posterior means for drift-diffusion parameters on day two.

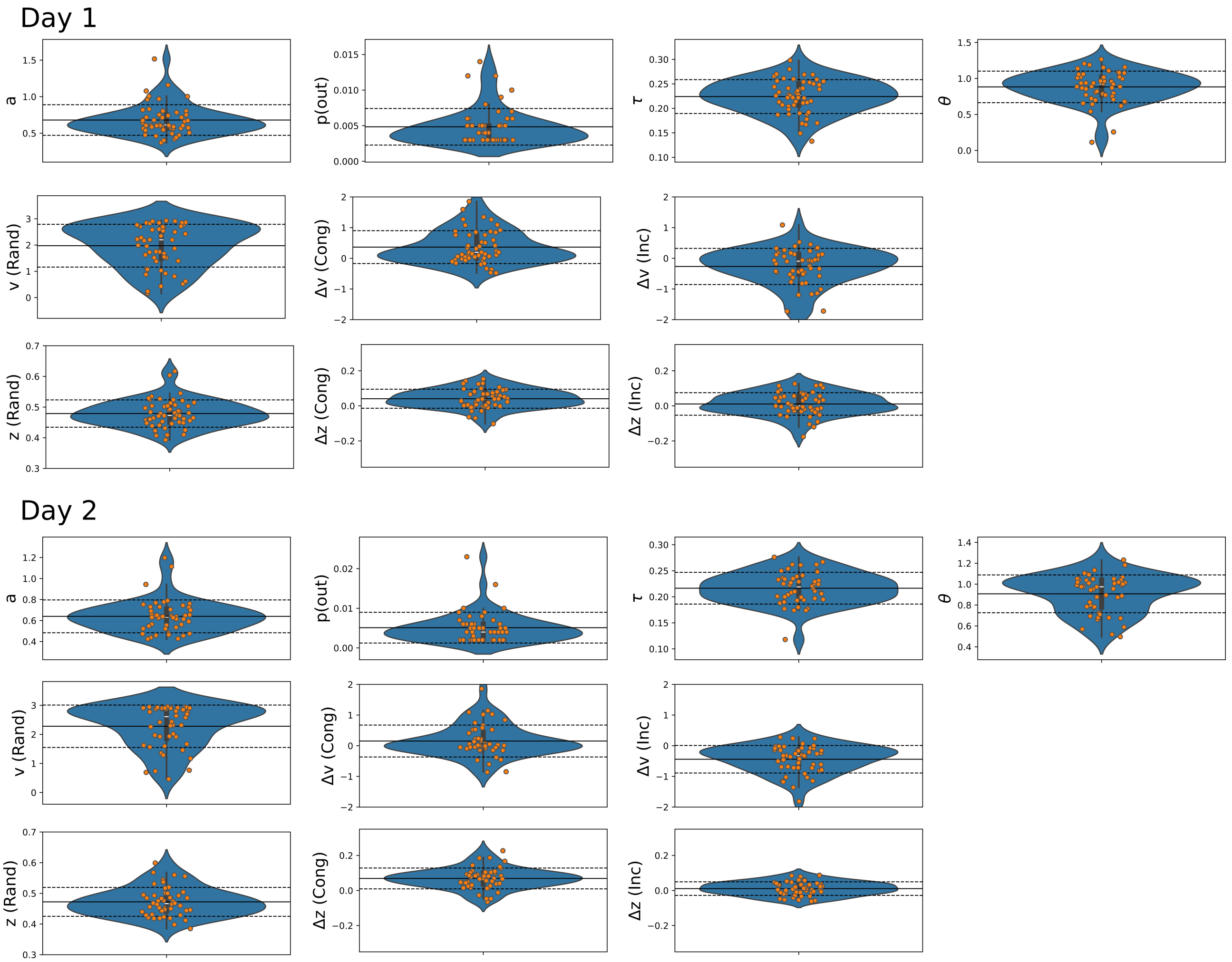


**Figure S5: DDM Results**: Means of posterior distributions for individual participants. *a* Boundary separation, *p(out)* probability of a random choice, $\tau$ non-decision time (in seconds), $\theta$ boundary angle (in radians), *v (Rand)* drift rate in random DTTs, $\Delta$ *(Cong)* Difference of drift rate in congruent DTTs compared to random DTTs, $\Delta$ *(Inc)* Difference of drift rate in incongruent DTTs compared to random DTTs, *z (Rand)* starting point bias in random DTTs, $\Delta$ *(Cong)* Difference of starting point bias in congruent DTTs compared to random DTTs, $\Delta$ *(Inc)* Difference of starting point bias in incongruent DTTs compared to random DTTs.

### iv. Post-Experiment Questionnaire

**Question 1: Did you have the impression that there were phases in which the experiment appeared easier?** Yes: 31 (51.67 %), No: 28 (46.67 %), Don't know: 1 (1.67 %).
**Question 2: Throughout the experiment, there were phases in which a sequence of 12 button presses was repeated often. did you notice this?** Yes: 16 (26.67 %), No: 36 (60 %), Don't know: 8 (13.34 %).
**Question 3: Throughout the experiment, there were phases in which a sequence of 12 button presses was repeated often. Please try to reproduce the sequence, or at least parts of it, by entering the corresponding keys (s,x,k,m) in the order of the sequence in the field below.** The longest correctly reproduced sub-sequence was on average $3.8 \pm 1.6$ elements long. No participant was able to reproduce more than 9 correct elements of the repeating action sequence.

### v. Effects of noticing the repeating action sequence and age effects

**Noticing the repeating action sequence.** Of 60 participants, 16 (26.67%) reported in a post-experiment questionnaire that they had noticed a repeating sequence in the experiment, 36 (60%) participants reported that they did not notice a sequence, and 8 (13.34%) participants answered "Don't know". No participant was able to reproduce the complete action sequence. T-tests between participants who did report that they noticed and those who did not notice a sequence revealed that noticing the sequence was associated with reduced optimal responding in random DTTs ($t(50) = -2.7, p = 0.01$), and with a significantly greater difference of optimal responding between congruent and random ($t(50) = 3.5, p = 0.001$), random and incongruent DTTs ($t(50) = 2.3, p = 0.03$), and congruent and incongruent context ($t(50) = 3.4, p = 0.001$).

**Age effects.** As expected, higher age correlated positively with reaction times, in both DTT ($r = 0.36, p = 0.004$) and STT ($r = 0.34, p = 0.007$). Age did not correlate with error rates ($p > 0.21$). Age did also not correlate with optimal responding in any of the three DTT types ($p > 0.32$). Importantly, measures of habit-learning, such as $\Delta RT$, $\Delta ER$, and the differences in optimal responses between the three dual-target trial types, did also not significantly correlate with age, nor show a trend. The correlation between ΔRT and the difference of optimal responses between the congruent and incongruent contexts ($C - I$) was higher in young participants than in old participants on both days, although the difference was not significant ($p = 0.1$) (median-split comparison between correlation values with two-tailed Z-test on Fisher-transformed correlation values). Participants who noticed a sequence were of similar age as those who did not (two-sample t-test: $t(50) = 1.0$, $p = 0.31$).
